# Deleterious genetic variants in *NOTCH1* are a major contributor to the incidence of non-syndromic Tetralogy of Fallot

**DOI:** 10.1101/300905

**Authors:** Donna J. Page, Matthieu J. Miossec, Simon G. Williams, Elisavet Fotiou, Richard M. Monaghan, Heather J. Cordell, Louise Sutcliffe, Ana Topf, Mathieu Bourgey, Guillaume Bourque, Robert Eveleigh, Sally L. Dunwoodie, David S. Winlaw, Shoumo Bhattacharya, Jeroen Breckpot, Koenraad Devriendt, Marc Gewillig, David Brook, Kerry Setchfield, Frances A. Bu’Lock, John O’Sullivan, Graham Stuart, Connie Bezzina, Barbara J.M. Mulder, Alex V. Postma, James R. Bentham, Martin Baron, Sanjeev S. Bhaskar, Graeme C. Black, William G. Newman, Kathryn E. Hentges, Mark Lathrop, Mauro Santibanez-Koref, Bernard D. Keavney

## Abstract

**Aims:** Familial recurrence studies provide strong evidence for a genetic component to the predisposition to sporadic, non-syndromic Tetralogy of Fallot (TOF), the most common cyanotic congenital heart disease (CHD) phenotype. Rare genetic variants have been identified as important contributors to the risk of CHD, but relatively small numbers of TOF cases have been studied to date. Here, we use whole exome sequencing to assess the prevalence of rare, potentially deleterious variants in candidate genes previously associated with both syndromic and non-syndromic TOF, in the largest cohort of non-syndromic TOF patients reported to date.

**Methods & Results:** 829 non-syndromic TOF patients underwent whole exome sequencing. A systematic review of the literature was conducted which revealed 77 genes in which mutations had been reported in patients with TOF. The presence of rare, deleterious variants in the 77 candidate genes was determined, defined by a minor allele frequency of ≤ 0.001 and scaled combined annotation-dependent depletion (CADD) score of ≥ 20. We found a clustering of heterozygous rare, deleterious variants in *NOTCH1* (P=1.89E-15), *DOCK6* (P=2.93E-07), *MYOM2* (P= 7.35E-05), *TTC37* (P=0.016), *MESP1* (P=0.024) and *TBX1* (P=0.039), after correcting for multiple testing. *NOTCH1* was most frequently found to harbour deleterious variants. Changes were observed in 49 patients (6%; 95% confidence interval [CI]: 4.5% - 7.8%) and included six truncating/frameshift variants and forty missense variants. Sanger sequencing of the unaffected parents of thirteen cases identified five *de novo* variants. Variants were not confined to a single functional domain of the NOTCH1 protein but significant clustering of variants was evident in the EGF-like repeats (P=0.018). Three *NOTCH1* missense variants (p.G200R, p.C607Y and *de novo* p.N1875S) were subjected to functional evaluation and showed a reduction in Jagged1 ligand-induced NOTCH signalling. p.C607Y, which exhibited the most significant reduction in signalling, also perturbed S1 cleavage of the NOTCH1 receptor in the Golgi.

**Conclusion:** The *NOTCH1* locus is a frequent site of genetic variants predisposing to non-syndromic TOF with 6% of patients exhibiting rare, deleterious variants. Our data supports the polygenic origin of TOF and suggests larger studies may identify additional loci.

## Introduction

Congenital heart disease (CHD) is the most common type of birth defect, affecting 8/1000 live births (1). CHD covers a large spectrum of heterogeneous cardiovascular phenotypes that range from single, localised defects to more complex structural abnormalities. Tetralogy of Fallot (TOF) is the most common complex, cyanotic CHD with a prevalence of 1/3000 births (1,2). TOF is considered a malformation of the cardiac outflow tract which comprises four specific structural characteristics postnatally; a ventricular septal defect (VSD), anterocephalad deviation of the outflow septum with resultant overriding of the aorta, variable obstruction of the right ventricular outflow tract (pulmonary stenosis) and consequent hypertrophy of the right ventricle (2,3). Surgical interventions during infancy mean that 85-90% of TOF patients now survive until at least 30 years of age (1,4). However, this is not without consequence; event-free survival is just 25% after 40 years of age (5) since resultant scar tissue from surgery and pulmonary regurgitation cause significant morbidity in adulthood (6,7).

The cause of TOF is elusive and no single candidate gene can be held accountable for the disease phenotype. However, the genetic status of syndromic TOF sufferers has provided valuable insights into causative genes in some patients. Approximately 20% of cases are associated with a recognised syndrome or chromosomal anomaly (2). Most significantly, approximately 15% of TOF patients have 22q11.2 deletion syndrome, wherein the major causal gene is *TBX1* (8,9). In approximately 80% of TOF cases which are non-syndromic there is generally not an identifiable cause, largely due to their non-Mendelian pattern of inheritance (10–13). Accordingly, a polygenic genetic architecture has been hypothesised and genome-wide approaches have been undertaken to provide insights into the complex genetic alterations responsible for TOF and other CHDs (11,13–18).

Whole exome sequencing (WES) has been used successfully to identify new CHD candidate genes (14,17,19,20). Many lines of evidence indicate a degree of phenotypic specificity of variants in particular genes. For example, the spectrum of phenotypes caused by 22q11.2 deletion or mutations in *TBX1* typically involves the outflow tract and great vessels, while Down’s syndrome or mutations in *NKX2.5* typically cause septal defects. To date, no WES study of CHD has included substantial numbers of any homogeneous phenotype, which should *a priori* have the highest power to identify causal variants.

Here, we present findings from WES of the largest cohort of non-syndromic TOF patients reported to date. First, we carried out a systematic review of the literature and identified 77 genes previously associated with either syndromic or non-syndromic TOF. We subsequently performed WES in 829 TOF probands and identified rare deleterious variants in the candidate genes. We sought evidence of pathological relevance for a subset of variants in the most significantly over-represented gene, based on the variants’ *de novo* occurrence and functional consequences in cellular models.

## Methods Summary

829 TOF probands were subjected to WES and rare (minor allele frequency [MAF] ≤ 0.001 in the Exome Aggregation Consortium [ExAC] database), deleterious (combined annotation-dependent depletion [CADD] score of ≥ 20) variants in 77 TOF candidate genes were assessed. Any variants observed in 1252 reference exome samples, that were analysed using the same approach as our case data, were eliminated. Clustering analysis was used to identify genes where significantly more variants were observed than expected. *De novo* variants were identified by Sanger sequencing of proband parent samples. Immunoblotting and luciferase assays were used to assess the expression and signalling activity of selected variants. Detailed methods can be found in the **Supplementary Data**.

## Results

### The *NOTCH1* locus most frequently harbours rare, deleterious variants in non-syndromic TOF patients

We assessed the incidence of rare, deleterious variants in 77 TOF candidate genes (Supplementary Table 1), for 829 non-syndromic TOF cases. Any variants observed in 1252 reference exomes were removed from consideration as a potential TOF susceptibility variant (Supplementary Table 2). Of the 77 genes considered, *NOTCH1* was the locus most frequently found to harbour a rare (ExAC MAF ≤ 0.001), deleterious (CADD ≥ 20) variant among TOF patients (table 1), with 49 probands exhibiting 46 *NOTCH1* variants, accounting for 6% of our TOF patient cohort (95% CI: 4.5% - 7.8%). The second most frequent locus was *DOCK6*, in which 30 probands harboured 26 rare, deleterious variants (table 1). The combined number of samples with *NOTCH1* or *DOCK6* variants was 76, highlighting the minimal overlap between probands with variants in these two genes (table 1). The statistical significance of these findings was assessed for each gene using clustering analysis which corrected for gene size; this confirmed a significant excess of rare, deleterious variants in six genes (figure 1); *NOTCH1* (P= 1.89E-15), *DOCK6* (P=2.93E-7), *MYOM2* (P= 7.35E-05), *TTC37* (P=0.016), *MESP1* (P=0.024) and *TBX1* (P=0.039). 126 TOF cases had a rare, deleterious variant in one or more of these genes, accounting for over 15% of our patient cohort. Clustering of variants identified in the remaining 71 candidate genes did not reach a corrected, 5% level of significance (Supplementary Table 3). When relaxing our stringent filters by removal of the CADD filter (Supplementary Table 4) and by increasing the MAF to ≤ 0.01 (Supplementary Table 5), *NOTCH1* continues to be the highest risk locus for non-syndromic TOF, with 8% of our cohort carrying *NOTCH1* variants.

**Table 1:**
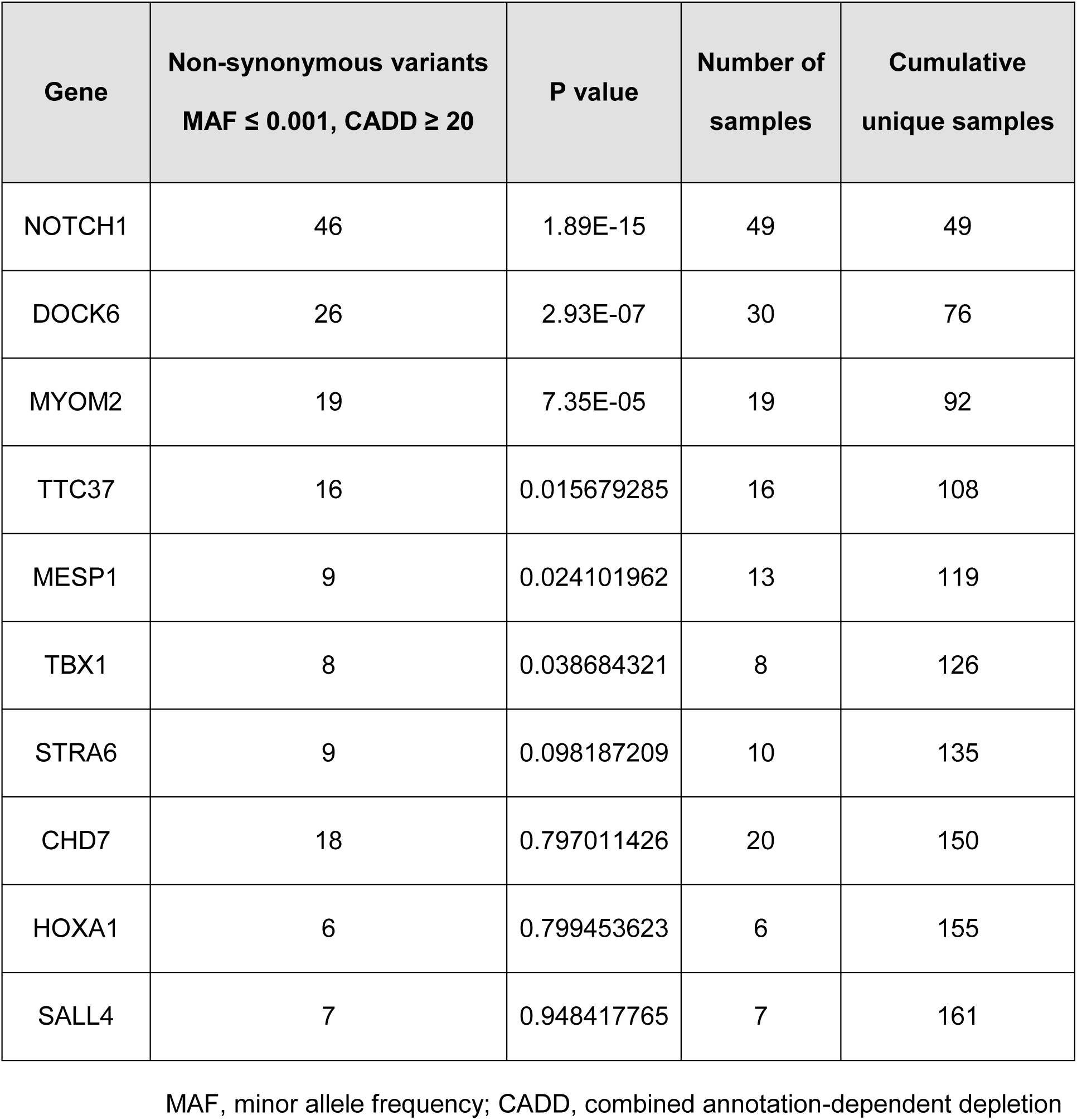
The top 10 genes ordered by levels of significance following the clustering analysis of rare, deleterious variants

| **Gene** | **Non-synonymous variants**<br>**MAF ≤ 0.001, CADD ≥ 20** | **P value** | **Number of**<br>**samples** | **Cumulative**<br>**unique samples** |
| --- | --- | --- | --- | --- |
| NOTCH1 | 46 | 1.89E-15 | 49 | 49 |
| DOCK6 | 26 | 2.93E-07 | 30 | 76 |
| MYOM2 | 19 | 7.35E-05 | 19 | 92 |
| TTC37 | 16 | 0.015679285 | 16 | 108 |
| MESP1 | 9 | 0.024101962 | 13 | 119 |
| TBX1 | 8 | 0.038684321 | 8 | 126 |
| STRA6 | 9 | 0.098187209 | 10 | 135 |
| CHD7 | 18 | 0.797011426 | 20 | 150 |
| HOXA1 | 6 | 0.799453623 | 6 | 155 |
| SALL4 | 7 | 0.948417765 | 7 | 161 |

MAF, minor allele frequency; CADD, combined annotation-dependent depletion

**Figure 1:**
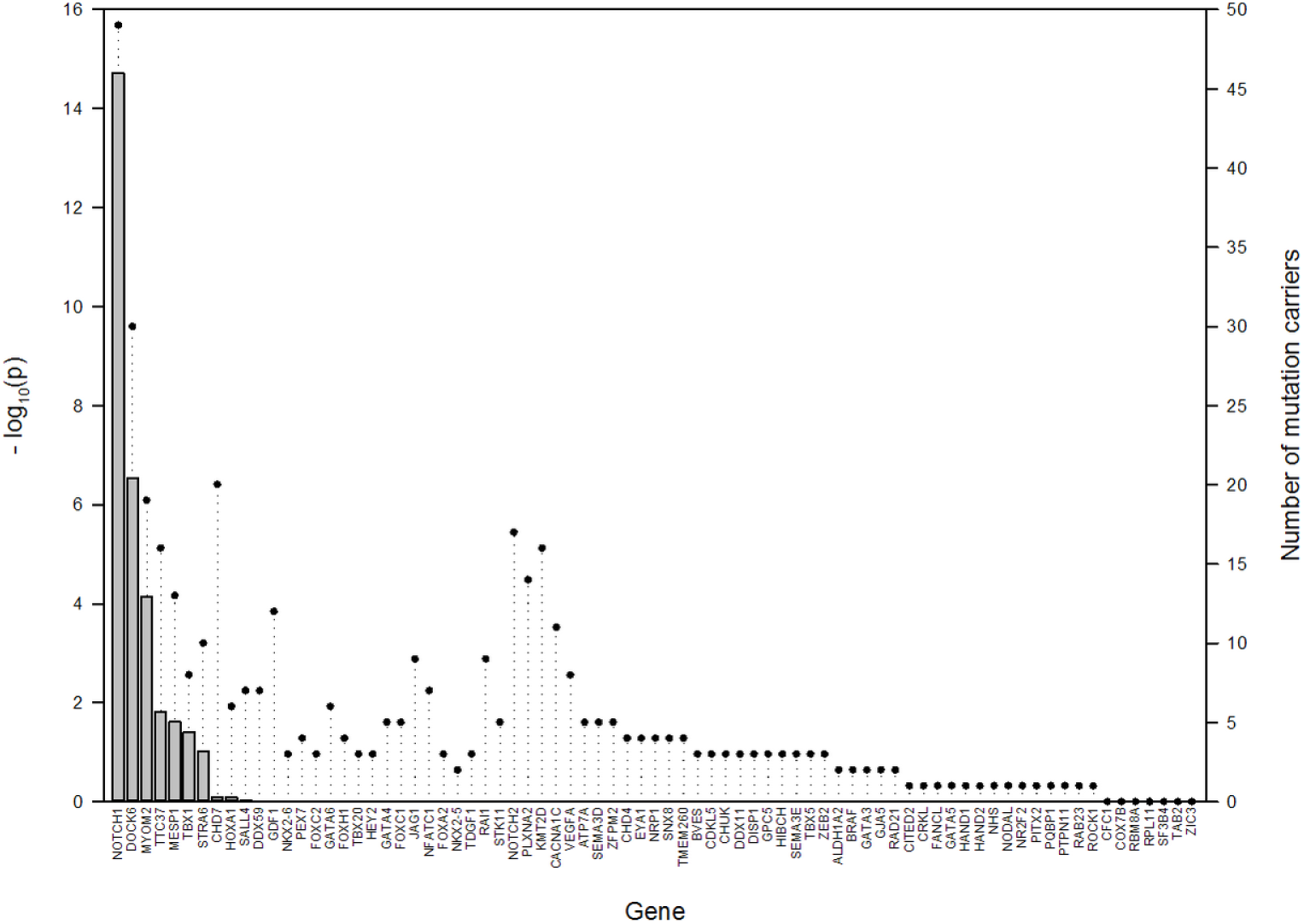
The number of patients carrying rare, deleterious variants in the 77 genes previously associated with syndromic or non-syndromic TOF. Bars indicate the respective significance levels of variant clustering for each gene, represented as –log P values. Circles represent the number of mutation carriers.

### *NOTCH1* variants cluster within the EGF-like repeats

We mapped the distribution of the 46 *NOTCH1* variants (Supplementary Table 6) to the various domains of the NOTCH1 protein (figure 2). Six of the variants identified were loss-of-function (LOF; figure 2), including three premature stop codons (p.R448^⋆^, p.W1638^⋆^ and p.Q1733^⋆^) and three single base pair deletions resulting in frameshifts and eventual premature truncation (p.G115fs^⋆^6, p.N147fs^⋆^128 and p.C1322fs^⋆^121). The three frameshift mutations were mapped to the EGF-like repeats in addition to one truncating mutant, p.R448^⋆^, whereas the remaining two truncating variants were located in the heterodimerisation domain (HD). Thus, all six LOF variants in *NOTCH1* were located in the extracellular domain of the protein. Of the remaining 40 missense variants, 21 were located in amino acid residues that are conserved with *Drosophila melanogaster* Notch. 22 of the missense variants were novel whereas the remaining 18 were reported in the ExAC database with very low frequency (MAF<0.001). Rare, deleterious variants were located throughout the NOTCH1 protein (figure 2), mapping to both the extracellular domain and the intracellular domain. However, significantly more variants were located in the EGF-like repeats (p=0.018; Supplementary Table 7), after adjustment for the proportion of the protein made up by EGF-like repeats. This region represents 56% of the total protein yet harbours 74% of *NOTCH1* variants. With appropriate caveat for this having been a *post hoc* analysis informed by our knowledge of the variant distribution, these data suggest that rare and deleterious variants in TOF are significantly enriched in the EGF-like repeats of NOTCH1. Of the intracellular domain mutants, two missense variants in the Ankyrin domain region, p.R2004L and p.A2036T, are particularly notable. R2004 is a surface exposed residue in Ankyrin domain 4 which is located in an interface region with the CSL transcription factor complex (21) and also located at an interface that binds the positive Notch regulator, Deltex (22). A2036 is located in Ankyrin domain 5 and lies immediately adjacent to the location of a *Drosophila* developmental mutant *N*^*Su*(*42C*)^ which is reported to reduce Notch activity (23).

**Figure 2:**
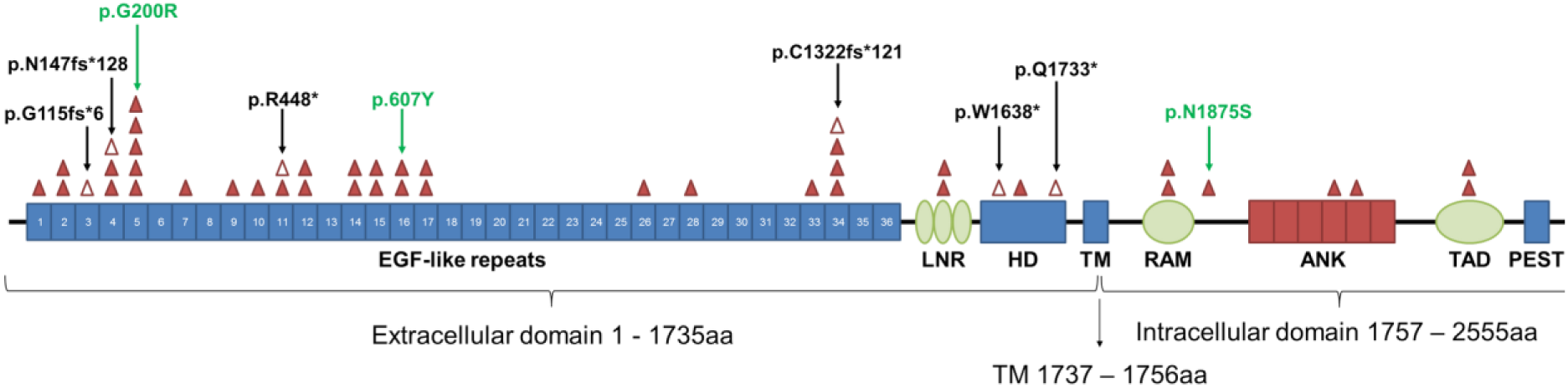
The NOTCH1 protein structure and location of rare, deleterious mutations in TOF patients. Variants were mapped to the known protein domains of NOTCH1. Missense variants are indicated by closed triangles and loss-of-function (truncating/frameshift) variants are indicated by open triangles and annotated in black. Missense variants p.G200R, p.C607Y and p.N1875S, which are of relevance to subsequent analyses, are indicated in green. ANK, ankyrin repeats; EGF, epidermal growth factor; HD, heterodimerisation domain; LBR, ligand binding region; LNR, Lin/Notch repeats; PEST, PEST domain; RAM, RBPJ-associated molecule domain; TAD, transactivation domain; TM, transmembrane domain.

### *De novo* variants identified in *NOTCH1*

We investigated the occurrence rate of *de novo* mutations in probands with *NOTCH1* variants. Of the 49 probands in our TOF patient cohort that harboured rare, deleterious variants in *NOTCH1*, samples from both parents were available for thirteen probands and analysed for variant inheritance. Following Sanger sequencing, five of the thirteen *NOTCH1* variants tested were identified as *de novo;* two of these were truncating variants, whereas the remaining three *de novo* variants were missense (table 2). All *de novo* variants identified were absent in the ExAC database and not previously reported in the literature. The remaining variants sequenced were inherited from parents and five of these were of maternal origin. Of the eight inherited variants, six were listed in ExAC with very low frequency (table 2); the remaining two variants have not been previously reported. These findings are in keeping with the results of previous WES experiments in CHD, where rare transmitted variants with strong bioinformatic support for functional impact, which are of presumed incomplete penetrance, have been uniformly encountered (14,17,20).

**Table 2:** Sequencing of parent samples to determine *NOTCH1* variant inheritance

| Amino acid change | Ref | Alt | LOF | Impact | In ExAC | MAF | Inheritance status |
| --- | --- | --- | --- | --- | --- | --- | --- |
| p.V53M | C | T | NO | Missense variant | YES | 3.69E-05 | FROM MOTHER |
| p.G200V | C | A | NO | Missense variant | NO | - | *DE NOVO* |
| p.C292Y | C | T | NO | Missense variant | NO | - | FROM MOTHER |
| p.P407L | G | A | NO | Missense variant | YES | 8.34E-06 | FROM MOTHER |
| p.T432M | G | A | NO | Missense variant | YES | 0.000157 | FROM FATHER |
| p.R448* | G | A | YES | Stop gained | NO | - | *DE NOVO* |
| p.T984M | G | A | NO | Missense variant | YES | 8.26E-06 | FROM MOTHER |
| p.Q1495K | G | T | NO | Missense variant | NO | - | FROM FATHER |
| p.C1549Y | C | T | NO | Missense variant | NO | - | *DE NOVO* |
| p.W1638* | C | T | YES | Stop gained | NO | - | *DE NOVO* |
| p.N1875S | T | C | NO | Missense variant | NO | - | *DE NOVO* |
| p.A2036T | C | T | NO | Missense variant | YES | 8.26E-06 | FROM FATHER |
| p.R2327Q | C | T | NO | Missense variant | YES | 0.000224 | FROM MOTHER |

Ref, reference allele; Alt, alternate allele; LOF, loss of function; MAF, minor allele frequency

### *NOTCH1* variants cluster within the EGF-like repeats

We mapped the distribution of the 46 *NOTCH1* variants (Supplementary Table 6) to the various domains of the NOTCH1 protein (figure 2). Six of the variants identified were loss-of-function (LOF; figure 2), including three premature stop codons (p.R448^⋆^, p.W1638^⋆^ and p.Q1733^⋆^) and three single base pair deletions resulting in frameshifts and eventual premature truncation (p.G115fs^⋆^6, p.N147fs^⋆^128 and p.C1322fs^⋆^121). The three frameshift mutations were mapped to the EGF-like repeats in addition to one truncating mutant, p.R448^⋆^, whereas the remaining two truncating variants were located in the heterodimerisation domain (HD). Thus, all six LOF variants in *NOTCH1* were located in the extracellular domain of the protein. Of the remaining 40 missense variants, 21 were located in amino acid residues that are conserved with *Drosophila melanogaster* Notch. 22 of the missense variants were novel whereas the remaining 18 were reported in the ExAC database with very low frequency (MAF<0.001). Rare, deleterious variants were located throughout the NOTCH1 protein (figure 2), mapping to both the extracellular domain and the intracellular domain. However, significantly more variants were located in the EGF-like repeats (p=0.018; Supplementary Table 7), after adjustment for the proportion of the protein made up by EGF-like repeats. This region represents 56% of the total protein yet harbours 74% of *NOTCH1* variants. With appropriate caveat for this having been a *post hoc* analysis informed by our knowledge of the variant distribution, these data suggest that rare and deleterious variants in TOF are significantly enriched in the EGF-like repeats of NOTCH1. Of the intracellular domain mutants, two missense variants in the Ankyrin domain region, p.R2004L and p.A2036T, are particularly notable. R2004 is a surface exposed residue in Ankyrin domain 4 which is located in an interface region with the CSL transcription factor complex (21) and also located at an interface that binds the positive Notch regulator, Deltex (22). A2036 is located in Ankyrin domain 5 and lies immediately adjacent to the location of a *Drosophila* developmental mutant *N*^*Su*(*42C*)^ which is reported to reduce Notch activity (23).

### *De novo* variants identified in *NOTCH1*

We investigated the occurrence rate of *de novo* mutations in probands with *NOTCH1* variants. Of the 49 probands in our TOF patient cohort that harboured rare, deleterious variants in *NOTCH1*, samples from both parents were available for thirteen probands and analysed for variant inheritance. Following Sanger sequencing, five of the thirteen *NOTCH1* variants tested were identified as *de novo;* two of these were truncating variants, whereas the remaining three *de novo* variants were missense (table 2). All *de novo* variants identified were absent in the ExAC database and not previously reported in the literature. The remaining variants sequenced were inherited from parents and five of these were of maternal origin. Of the eight inherited variants, six were listed in ExAC with very low frequency (table 2); the remaining two variants have not been previously reported. These findings are in keeping with the results of previous WES experiments in CHD, where rare transmitted variants with strong bioinformatic support for functional impact, which are of presumed incomplete penetrance, have been uniformly encountered (14,17,20).

### *NOTCH1* variants can affect NOTCH1 processing and signalling function

The *NOTCH1* gene encodes an evolutionarily conserved transmembrane receptor that mediates cell-cell communication to govern cell fate decisions during development (24). S1 cleavage is an important step in the maturation of the NOTCH1 receptor. During this process, the 300 kDa translation product of NOTCH1 undergoes cleavage in the Golgi by furin-like convertase to generate two polypeptides of 180 and 120 kDa (25). To determine whether *NOTCH1* variants affect S1 cleavage, we assessed the expression of three NOTCH1 variants in comparison to WT NOTCH1 by immunoblotting. The variants assessed were p.G200R, p.C607Y and p.N1875S (figure 2); p.G200R is located in a conserved residue located within a β-hairpin turn within EGF5, and p.C607Y, located in EGF16, removes a conserved disulphide bond that normally would be expected to stabilise the EGF-domain conformation. p.N1875S is located in a residue that lies in a linker region between the RAM and Ankyrin regions of the Notch intracellular domain. As expected, we observed two bands at 300 kDa (P300) and 120 kDa (P120), representing full length and cleaved NOTCH1 protein (25); the remaining 180 kDa product was not detectable due to the positioning of our FLAG-tag at the C-terminus (figure 3a). For WT NOTCH1, p.G200R and p.N1875S variants, we observe similar levels of both P300 and P120 (figure 3a). However, the p.C607Y variant exhibited perturbed cleavage by furin-like convertase (25). Indeed, quantification confirmed that 5%±0.37% of NOTCH1 p.C607Y underwent cleavage in comparison to 57%±3.96% of WT NOTCH1 (P=0.0002; figure 3b). Hence, the p.C607Y variant affects the processing of NOTCH1, whereas the receptor is processed normally in the presence of p.G200R and p.N1875S variants.

**Figure 3:**
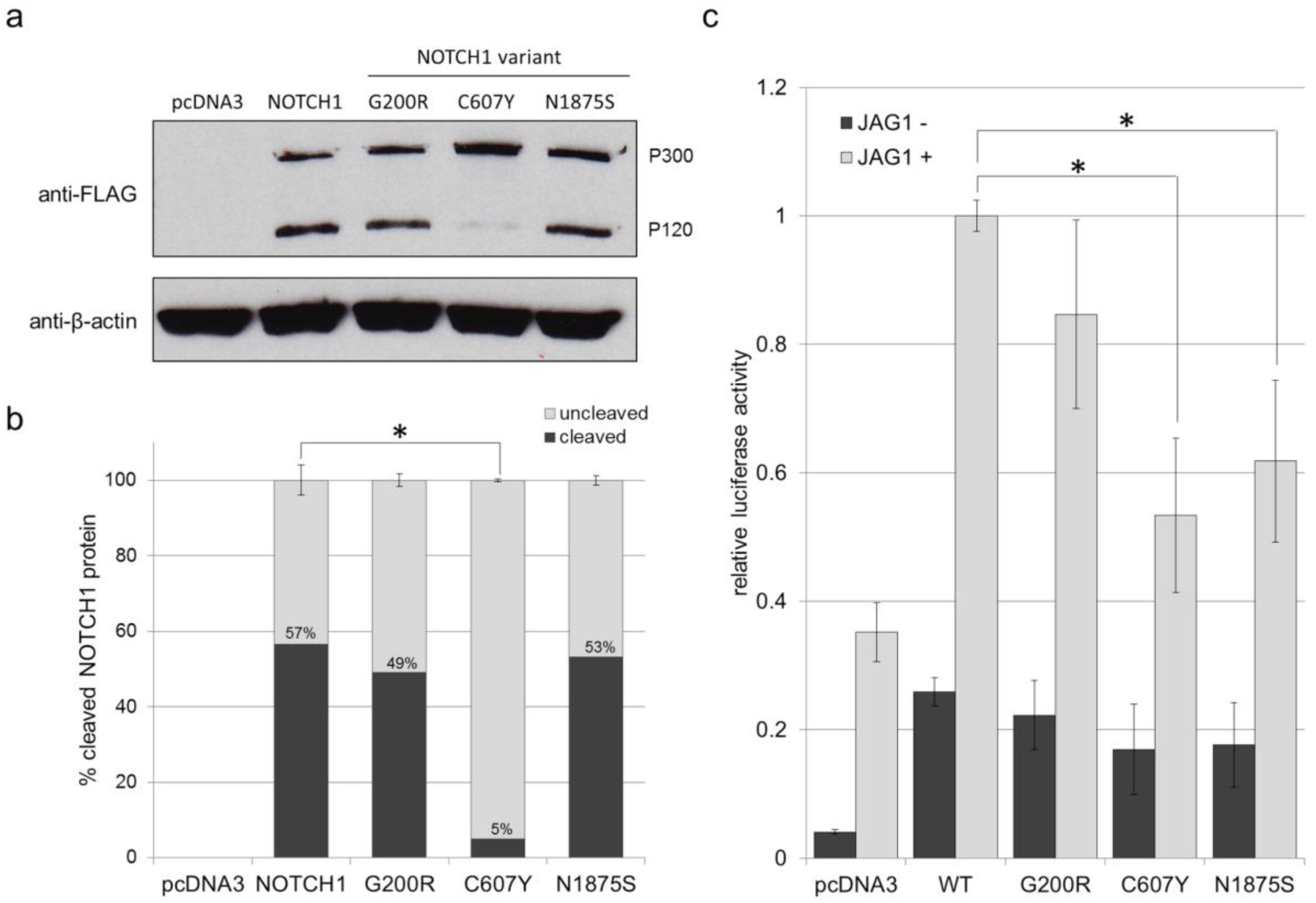
(a) Immunoblot for FLAG to determine the expression and cleavage of NOTCH1 variants p.G200R, p.C607Y and p.N1875S in comparison to WT NOTCH1 following overexpression in HeLa cells. The two bands at 300 kDa (P300) and 120 kDa (P120) represent the full length and cleaved NOTCH1 protein. β-actin was used as a loading control. (b) Quantification of the percentage of cleaved versus uncleaved NOTCH1 protein for WT NOTCH1 and NOTCH variants p.G200R, p.C607Y and p.N1875S. Error bars: mean ±SEM from three biological replicates and statistical significance was determined using two-tailed paired *t*-tests. (c) The effect of rare, deleterious *NOTCH1* variants on Jagged-induced NOTCH signalling levels. NOTCH signaling activity was measured using a luciferase-based reporter system (RBPJ). HeLa cells were cultured with or without immobilised JAG1 ligand and co-transfected with RBPJ reporter constructs and WT NOTCH1, p.G200R, p.C607Y or p.N1875S. Firefly luciferase readings were normalised to Renilla luciferase readings to control for transfection efficiency and cell number. RBPJ activity was expressed relative to WT for comparison. Error bars: mean ±SEM from four biological replicates, each with three technical replicates. Statistical significance was assessed using two-tailed paired *t*-tests.

Heterodimeric NOTCH1 is membrane tethered and undergoes further cleavage by γ-secretase which releases the NOTCH intracellular domain (NICD). NICD subsequently translocates to the nucleus where it interacts with transcription factor RBPJ to activate NOTCH target genes (24). To determine whether p.G200R, p.C607Y and p.N1875S variants affect NOTCH1 canonical signalling function, we assessed NOTCH signalling through the canonical, RBPJ transcription factor-dependent pathway following stimulation with immobilised Jagged1 ligand. The variants were overexpressed in HeLa cells and NOTCH1 signalling was assessed by RBPJ luciferase activity. All three variants demonstrated reduced NOTCH signalling via RBPJ (figure 3c). The p.C607Y variant, that exhibited perturbed cleavage, significantly reduced NOTCH signalling by 47%±0.12% (P=0.008) compared to WT NOTCH1. Similarly, *de novo* variant p.N1875S reduced NOTCH signalling by 38%±0.13% (P=0.02). The p.G200R variant reduced NOTCH signalling by 15%±0.15%, although this finding was not significant (P=0.33) (figure 3c). No significant differences were observed between WT NOTCH1, p.G200R, p.C607Y and p.N1875S variants in the absence of JAG1 ligand. In each transfection experiment, mRNA expression of WT NOTCH1 and the three NOTCH1 variants was equal (data not shown), thus the differences in NOTCH1 signalling observed were not due to reduced mRNA expression of the variants. Hence, all three variants identified in patients that were subjected to functional testing were shown to affect canonical NOTCH1 signalling.

## Discussion

TOF is the most common, severe cyanotic CHD; however, variants that could account for the high degree of genetic susceptibility inferred from familial recurrence risk studies (26) are as yet unidentified. Indeed, no single gene locus, with the exception of the 22q11 deletion, has been found to account for any significant proportion of TOF cases. Through WES of a large cohort of sporadic, non-syndromic TOF, we show that *NOTCH1* is an important susceptibility gene; 6% of patients carry heterozygous variants in *NOTCH1* which, based on ExAC allele frequency, bioinformatic *in silico* prediction, and functional characterisation, we judged to be likely susceptibility alleles. Of the variants identified, 57% were novel, and the remainder have previously been reported in ExAC, but with very low frequency. Six of the variants were LOF, including truncating and frame shift mutations, whereas the remaining forty variants were missense and anticipated to be pathogenic. Five out of thirteen variants tested were *de novo*, adding to the evidence for pathogenicity; however the remaining variants were transmitted from unaffected parents indicating incomplete penetrance. As an additional safeguard against false positive results due to systematic methodological differences between our cohort and the studies which contributed to the ExAC database, we studied a set of over 1000 reference exomes in patients free from CHD, generated and analysed in the same fashion as the case exomes, removing any variant that appeared even once in the reference exome set from consideration as a potential TOF susceptibility variant.

Previous sequencing studies of CHD have identified an association of *NOTCH1* variants in left-sided cardiac malformations including bicuspid aortic valve, aortic valve stenosis, coarctation of the aorta and hypoplastic left heart syndrome (27–30). Like TOF, the causative *NOTCH1* variants described for these conditions are mostly missense with a small number of LOF variants located throughout the NOTCH1 protein. In contrast, few studies, which included only a small number of patients, have implicated *NOTCH1* variants in non-syndromic TOF (27,31). However, there are no clear distinctions between the type and location of *NOTCH1* variants identified in TOF compared to those reported in other isolated cardiovascular abnormalities. We therefore propose that genetic background and/or environmental influences may specify phenotypic expressivity.

The association of *NOTCH1* with cardiac defects beyond left-sided lesions is consistent with the reported roles of NOTCH1 during heart development. Active NOTCH1 is observed in trabecular endocardium and both global and endothelial-specific knockout of *Notch1* in mice results in abnormal ventricular trabeculae and abnormal cardiomyocyte patterning (32). Relevant to TOF, Notch1 plays a role in the organisation of the outflow tract, which requires the specification of cells from both the neural crest and secondary heart field (33). Furthermore, Notch1 is important for endocardial epithelial-to-mesenchymal transition, a process that is essential for cardiac valve formation (29,34). It should however be noted that all *NOTCH1* variants we report are heterozygous. There are numerous reports of global and tissue specific *NOTCH1* heterozygous mice that appear phenotypically normal, with no obvious cardiovascular pathologies (35,36). However, in a more recent study that assessed ascending aortic aneurysm, *NOTCH1*^+/-^ in a predominantly 129S6 background developed aortic root dilation; and this was in contrast to *NOTCH1*^+/-^ in a mixed background (37). These findings highlight the importance of genetic background in disease expressivity and are consistent with the incomplete penetrance observed.

Mutations in key cardiac transcription factors such as *NKX2.5* (38), *GATA4* (39), *HAND2* (12) and *GATA6* (40) have been identified in TOF cases, typically by targeted candidate gene sequencing; however these genes appear to account for just 1.7% of cases in our cohort. A Genome Wide Association Study (GWAS) of TOF cases versus controls identified risk alleles for single nucleotide polymorphisms (SNPs) in chromosomal regions 12q24 and 13q32, including the *PTPN11* and *GPC5* loci, respectively (13,16). Additionally, duplications in 1q21.1 are strongly enriched in TOF patients (11,31). Our findings, which in addition to *NOTCH1* showed significant enrichment in five other genes (including *TBX1*, a well-established TOF risk gene which is principally responsible for the cardiac manifestations of 22q11 deletion) concur with previous studies regarding the marked locus heterogeneity of the condition. A possible role for *NOTCH1* in non-syndromic TOF has previously been suggested by CNV analysis. A study of 34 infants with non-syndromic TOF revealed two patients with CNVs encompassing the *NOTCH1* gene (41). Additionally, a microdeletion including the *NOTCH1* locus in a patient with TOF was identified in a study of CNVs in 114 TOF patients (31). A recent study that focused primarily on families with left-sided CHD, identified family members with TOF harbouring pathogenic mutations in *NOTCH1* (27). Further indirect evidence came from a study that analysed the gene expression patterns in TOF patient right ventricles and found many genes from the *NOTCH* and *WNT* signalling pathways were significantly reduced. Interestingly, down-regulation of *NOTCH* signalling components was also observed in TOF patients with a 22q11.2 deletion (42), highlighting a common transcriptional signature between both syndromic and non-syndromic TOF, initiated by different genetic events. However, the present study involves, by a substantial margin, the largest TOF cohort studied by WES to date, providing the most accurate estimate thus far of the contribution of *NOTCH1* variants to TOF risk.

*De novo* mutations are a significant cause of early-onset genetic disorders, including CHD. Of the *NOTCH1* variants identified in this study, where parents were available, five of thirteen variants were found to be *de novo*, all of which were novel variants not previously reported on ExAC. Furthermore, *de novo* variant p.N1875S was shown to have significantly reduced Jagged1-induced NOTCH signalling relative to WT NOTCH1, providing further support as to the pathogenicity of *de novo* variants. However, other variants were inherited from an unaffected parent, confirming the role of incompletely penetrant variants observed for other CHD genes and phenotypes (17,20). The incomplete penetrance is in keeping with the complex genetic aetiology of non-syndromic TOF, in which families segregating the condition in a Mendelian fashion are rarely encountered and genetic background in addition to *in utero* environmental factors can be inferred to play significant roles.

Recently, Blue *et al* (2014) identified the *NOTCH1* missense variant p.G200R, which was independently found in the present study, to segregate with disease in two cousins with right-sided CHD. Cardiovascular malformations included persistent truncus arteriosus, VSD, pulmonary atresia, and major aorto-pulmonary collateral arteries. Furthermore, a case of TOF was also reported in the preceding generation, although sequencing analysis was not carried out on this relative. Our functional assessment of this variant showed reduced Jagged1-induced canonical NOTCH signalling. *NOTCH1* is extremely LOF intolerant, with a probability of being LOF intolerant (pLI) of 1.0 on ExAC. Thus, even mild changes in its signalling ability could potentially have negative consequences during development. Hence, co-segregation of p.G200R with disease, in addition to reduced signalling capacity in *in vitro* experimentation provide evidence as to possible pathogenicity of this variant, albeit with incomplete penetrance (43).The p.C607Y missense variant perturbed NOTCH1 receptor cleavage by the calcium-dependent enzyme, furin-like convertase. The cleavage site is located at amino acids 1651 - 1654, some distance away from the variant. A similar observation has been reported by McBride *et al* (2008) where *NOTCH1* variant p.A683T, identified in two patients with left ventricular outflow tract malformations, also perturbed S1 cleavage by similar levels. In both cases, this led to a 50% reduction in RBPJ luciferase activity (44). The mechanism by which such variants alter S1 cleavage requires further research.

Autosomal dominant germ-line mutations in the *NOTCH1* gene are one of the causes of Adams-Oliver syndrome (AOS) which is chiefly characterised by *aplasia cutis congenita* and terminal transverse limb defects. In addition, around half of patients exhibit congenital cardiac anomalies, including atrial septal defect (ASD), VSD, aortic valve stenosis, pulmonary valve stenosis and TOF (51,52). AOS is an extremely rare syndrome, with a prevalence of approximately 1 in 225,000 (52). No patient in our cohort had diagnostic features of AOS. Like TOF, the causative *NOTCH1* variants described for AOS are both missense and LOF variants that are not confined to one specific protein domain; however, the majority of reported variants are similarly located in the EGF-like repeats (51,52). There are no clear distinctions between the *NOTCH1* variants we have identified in TOF versus those that cause AOS, though no previously described AOS variant was present in our cases. The extra-cardiac features of AOS have been suggested to be due to early embryonic vascular abnormalities (53); this raises the possibility that AOS, TOF and other cardiac anomalies that occur due to mutations in *NOTCH1* may be a spectrum of disorders. Other examples of syndromic genes that can cause isolated CHD, including TOF, are *PTPN11* (Noonan’s syndrome), (13,54), *TBX5* (Holt-Oram syndrome) (55) and *JAG1* (Alagille syndrome) (56). Determining the role of genetic background, environmental context and the specific *NOTCH1* variants in determining the severity of the cardiac phenotype and the occurrence of extra-cardiac malformations requires further research.

The second gene identified in the current study, *DOCK6*, has also been identified as a causative gene in AOS, responsible for an autosomal-recessive form, often associated with impaired vascular function (57,58). The *DOCK6* gene encodes a highly conserved member of the Dedicator of Cytokinesis family and functions as an atypical nucleotide exchange factor. Specifically, DOCK6 plays a role in the remodelling of the actin cytoskeleton by activating two members of the Rho GTPase family, CDC42 and RAC1 (59). Studies thus far have demonstrated a role for DOCK6 in axonal extension and branching during neuronal development (59,60). Presently, there is no simple functional read out for DOCK6, precluding detailed characterisation of the variants we discovered in cellular systems. However, investigations into a possible role for DOCK6 in cardiovascular development would be of interest.

In addition to *NOTCH1* and *DOCK6*, significant clustering of rare, deleterious variants was also observed in an additional four genes, *MYOM2*, *TTC37*, *MESP1 and TBX1*, accounting for approximately 4% of TOF cases in our cohort. These findings emphasise the genetic heterogeneity that may ultimately lead to a TOF phenotype. The *MYOM2* gene encodes Myomesin-2, a well characterised sarcomeric protein that is a major structural component of the myofibrillar M band. Mutations in *MYOM2* have been identified in hypertrophic cardiomyopathy (61) and more recently, Grunert *et al* (2014) reported four non-syndromic TOF cases with deleterious variants in *MYOM2. TTC37* encodes Tetratricopeptide Repeat Domain 37 responsible for Trichohepatoenteric syndrome 1 (THES). THES is an autosomal-recessive disorder characterised by life threatening diarrhoea during infancy accompanied by immunodeficiency, liver disease, facial dysmorphia, hypopigmentation and cardiac defects. Cardiac defects are reported in approximately 25% of THES cases and include TOF, VSD and ASD (62,63). The molecular function of *TTC37* is poorly characterised and its role during heart development unknown. *MESP1* encodes for Mesoderm posterior basic helix-loop-helix transcription factor 1, which plays an essential role during heart development. *Mesp1* is expressed in cardiac progenitors and participates in the timely specification of the cardiac lineage in mouse cardiac development (64–66). Investigations in mice suggest Mesp1 may act upstream of critical cardiac transcription factors including *Hand2*, *Gata4*, *Nkx2.5* and *Tbx20* (64,65). In a previous study, six rare, non-synonymous *MESP1* variants were identified in 647 patients with congenital conotruncal and related heart defects, including TOF (67). Finally, *TBX1* encodes T-box transcription factor 1 which has important roles during development including the migration of cardiac neural crest cells and cellular proliferation in the second heart field (68). *TBX1* is also responsible for the cardiac abnormalities associated with the 22q.11.2 deletion syndrome (69), which is the major contributor to the incidence of TOF (8,9). In agreement with Griffin *et al* (2010), the present study confirms that rare, deleterious *TBX1* variants contribute to a small proportion of non-syndromic TOF cases (70).

In summary, among the genes that have been implicated in TOF thus far, our large study indicates that of the 77 TOF candidate genes tested, no gene accounts for more than 8% (upper 95% binomial confidence interval) of non-syndromic TOF. Among these genes *NOTCH1* is the most commonly involved. The two most commonly involved genes (*NOTCH1* and *DOCK6*) are also both involved in the predisposition to AOS, suggesting further investigation of common pathways between these conditions may be fruitful. Some mutations in *NOTCH1* that are associated with TOF were *de novo*, but others were present in apparently asymptomatic individuals, indicating incomplete penetrance. Such incomplete penetrance has been prominently observed, for example, in Mendelian aortopathies, emphasising the importance of genetic background in structural cardiac and vascular diseases. Detailed phenotypic studies of mutation carriers who do not have overt CHD using advanced imaging may be of interest to delineate quantitative phenotypes potentially relevant to CHD.

## Acknowledgements

This work was supported by the British Heart Foundation. BK and SB hold BHF Personal Chairs. SB was supported by the BHF funded GOCHD study project grant. BM, CRB and AP were supported by the Netherlands Heart Foundation CVON project CONCOR-genes (CVON 2014-18). The work in Nottingham/Leicester was funded by BHF Programme Grant RG/13/10/30376. This study makes use of the ICR1000 UK exome series data generated by Professor Nazneen Rahman’s Team at The Institute of Cancer Research, London (71).

